# Exploring adaptation routes to low temperatures in the *Saccharomyces* genus

**DOI:** 10.1101/2024.02.25.582014

**Authors:** Javier Pinto, Laura Natalia Balarezo-Cisneros, Daniela Delneri

**Affiliations:** Manchester Institute of Biotechnology, Faculty of Biology Medicine and Health, The University of Manchester, Manchester, United Kingdom

## Abstract

The identification of traits that affect adaptation of microbial species to external abiotic factors, such as temperature, is key for our understanding of how biodiversity originates and can be maintained in a constantly changing environment. The *Saccharomyces* genus, which includes eight species with different thermotolerant profiles, represent an ideal experimental platform to study the impact of adaptive alleles in different genetic backgrounds. Previous studies identified a group of genes important for maintenance of growth at lower temperatures. Here, we carried out a genus-wide functional analysis in all eight *Saccharomyces* species for six candidate genes. We showed that the cold tolerance trait of *S. kudriavzevii* and *S. eubayanus* is likely to be evolved from different routes, involving genes important for the conservation of redox-balance, and for the long-chain fatty acid metabolism, respectively. For several loci, temperature- and species-dependent epistasis was detected, underlying the plasticity and complexity of the genetic interactions. The natural isolates of *S. kudriavzevii, S. jurei* and *S. mikatae* had a significantly higher expression of the genes involved in the redox balance compared to *S. cerevisiae*, raising the question of what proportion of the trait is accounted for solely due to transcriptional strength. To tease apart the role of gene expression from that of allelic variation, for two genes we independently replaced in four yeast species either the promoters or the alleles with those derived from *S. kudriavzevii*. Our data consistently showed a significant fitness improvement at cold temperatures in the strains carrying the *S. kudriavzevii* promoter, while growth was lower upon allele swapping. These results suggest that transcriptional strength plays a bigger role in growth maintenance at cold over the allele type and supports a model of adaptation centred on stochastic tuning of the expression network.

**Author summary:** The decline in biodiversity due to environmental changes influences the stability of ecosystems by altering the geographic distribution of several microbial and fungal species. Temperature is one of the leading factors that drive adaptation and different organisms share the same habitat because of their different thermal profiles. It is therefore important to study the genes that affect the fitness of microorganisms at different temperatures in order to understand both how biodiversity originated and how can be maintained. The *Saccharomyces* genus, which includes species with different thermotolerant profiles, represent an ideal experimental platform to investigate the impact of adaptive alleles in response to temperature changes. Here, we carried out a functional analysis for putative cold-tolerant genes and showed that this trait is likely to be evolved from different routes in different species, involving the conservation of redox-balance and alteration of membrane fluidity. Furthermore, for several species, genetic interactions display fitness tradeoffs in different environments. Finally, by unravelling the interplay between gene expression, allele variation, genetic background and environment, this study shed light on the intricate nature of transcriptional regulation and its pivotal role in facilitating cold adaptation in *Saccharomyces* species.

## Introduction

The fingerprint that human actions have left on the earth’s temperature has driven decline in biodiversity and influenced the stability of ecosystems by altering the geographic distribution of several species [1,2], including temperature sensitive microorganisms such as *Saccharomyces* yeast [3]. It is therefore important to understand the molecular mechanisms that affect the adaption and biodiversity of microbial species at different temperatures and how biodiversity originates and is maintained in a constantly changing environment [4,5,6,7]. Organisms can slowly adapt to a new environment by accumulating beneficial mutations in key genes or by rewiring parts of the regulatory networks hence changing gene expression [8].

The *Saccharomyces* genus is an ideal group to study interspecific ecological traits, including temperature, since it consists of a monophyletic group of species with high levels of sequence similarity within the *Ascomycota* phylum [9]. *Saccharomyces* consists of eight species that have been evolved and adapted to grow at a different range of temperatures; *S. kudriavzevii* (*Sku*), *S. arboricola* (*Sar*)*, S. uvarum* (*Suv*), *S. eubayanus* (*Seu*) and the novel species *S. jurei* (*Sju*) are considered cold-tolerant, *S. paradoxus* (*Spa*) is classified as a thermo-generalist (growing well on a broader range of temperatures), and finally *S. cerevisiae* (*Sce*) and *S. mikatae* (*Smi*) are more thermo-tolerant [10,11,12,13,14,15,16]. Studies to identify genes involved in temperature adaptation in wild *Saccharomyces* strains and species have been carried out over the last ten years [3,17,18]. For examples, a systems biology approach, coupling thermodynamic modelling with large-scale competition studies on the *S. cerevisiae* deletion collection, proved to be a valuable tool to identify a set of cold-tolerant genes, some of which were validated in two species with different thermoprofiles [18]. More recently, a set of thermo-tolerant genes is *S. cerevisiae* were introduced to the sibling species *S. paradoxus* and were shown to increase thermotolerance in this species by 15% [19]. Mitochondria also play a significant role in maintaining fitness at low temperatures in hybrid yeast species [20,21,22] and can influence nuclear transcription [23]. It has been shown that during temperature shifts the transcriptional network in hybrids show allelic bias and there is an overdominance of one ortholog set of parental alleles over the other [24]. Finally, functional analysis studies of non-coding RNAs (ncRNAs) in *S. cerevisiae* [25,26] have identify ncRNAs that influence gene transcription and growth at low temperature [27].

In this study, we investigate the impact of six non-essential candidate genes, identified as important for growth maintenance at cold in a large-scale study [18], in the eight *Saccharomyces* species, including *S. jurei*, a newly discovered *Saccharomyces* species from high altitude oaks [11]. The candidate genes are involved in a variety of cellular mechanisms and metabolic pathways that may affect the resistance to low temperatures, including synthesis of ethanol (*ADH3; ADH5*), glycerol utilisation (*GUT2*), NAD^+^ biosynthesis (*NMA1*), inhibition of glycotransferases (*YND1*), and fatty acid activation (*FAA1*) [28,29,30,31].

By analysing the impact of these genes on the fitness and gene expression within the *Saccharomyces* genus, we were able to identify species-dependent adaptation routes and temperature-dependant epistatic interactions. Experiments on both promoter and allele swap of *ADH3* and *YND1* between the cold-tolerant *S. kudriavzevii* and other four species revealed the main role of transcription over allele type in cold temperature adaptation.

Overall, our data show that cold tolerance can be enhanced in thermotolerant yeasts by altering transcription of specific genes and support the notion of stochastic transcription as selectable trait during adaptation to novel niches.

## Results

### Temperature-dependent growth profiling of *Saccharomyces jurei* and comparative analysis with other *Saccharomyces* species

Optimal growth temperatures and temperature ranges in which yeast isolates can grow and be maintained in the wild has been gathered over the last decade [32,33,34], however, no data are yet available for the newly discovered species *S. jurei*.

We determined the optimal growth temperature range for two *S. jurei* strains and conducted a comparative analysis across all the other *Saccharomyces* species. The *S. jurei* strains NCYC3947 and NCYC3962, showed an optimal growth temperature of 27.8°C and 27.2°C respectively, positioning them in temperature range between the more warm and more cold-tolerant species of the *Saccharomyces* genus, (Fig. 1). As expected, *S. cerevisiae* strain 96.2 had the highest optimal growth temperature (34.5°C) and *S. kudriavzevii* CAIII the lowest (23.6°C) (Fig. 1), confirming previous literature data (18,13,34). This data supports the idea that different preferential growing temperatures contributed to the diversification of *Saccharomyces* species, allowing them to share ecological niches.

**Figure 1.**
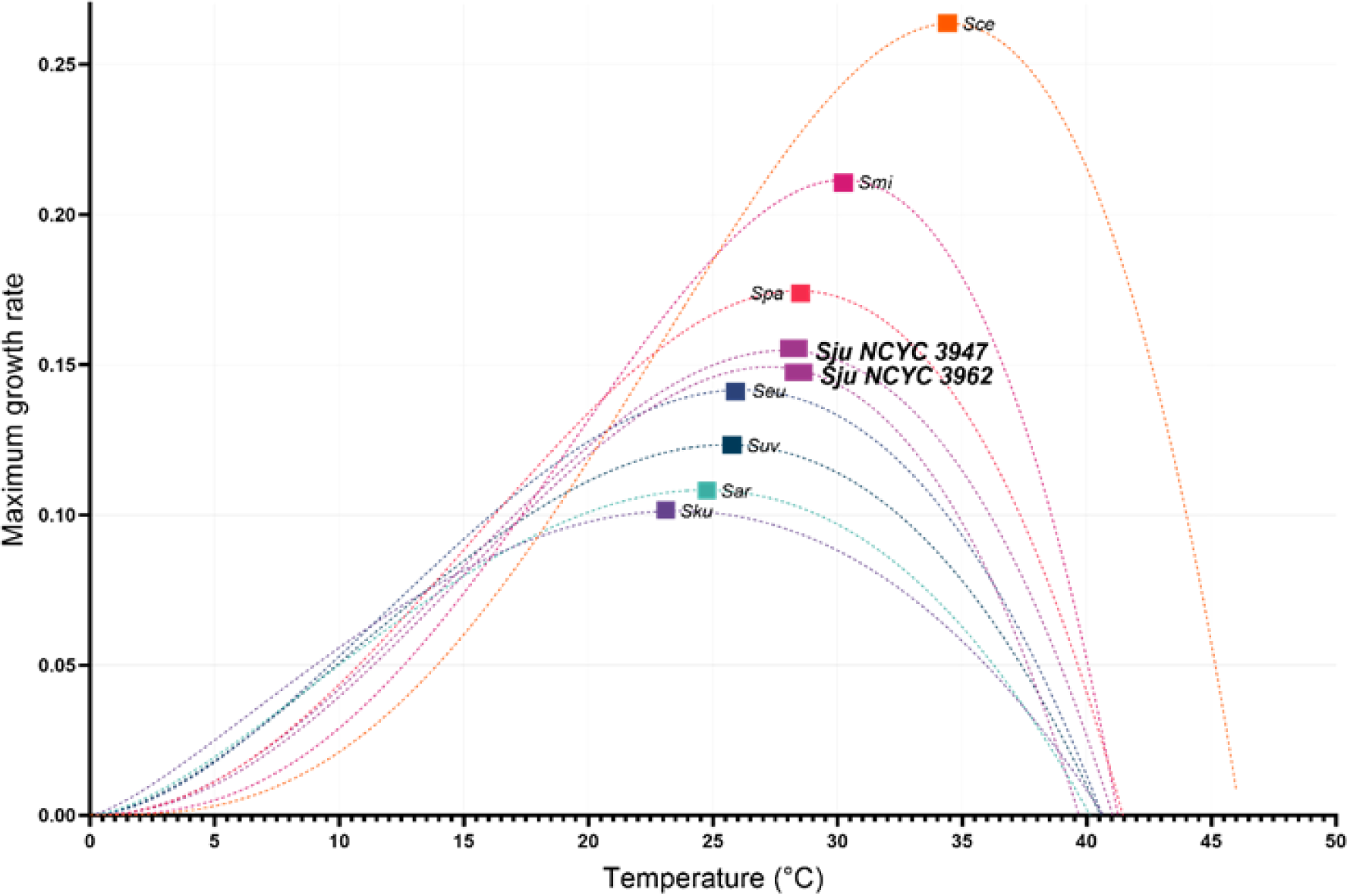
Maximum growth rate of the *Saccharomyces* species as a function of temperature. The figure includes *S. cerevisiae* (Sce), *S. paradoxus* (Spa), *S. mikatae* (Smi), *S. jurei* (Sju NCYC 3947), *S. jurei* (Sju NCYC 3962), *S. kudriavzevii* (Sku), *S. arboricola* (Sar), *S. eubayanus* (Seu) and *S. uvarum* (Suv) as a function of temperature. The graph was built using a non-lineal model based in observed fitness data obtained at 10°C, 15°C, 20°C, 25°C, 30°C, and 40°C.

### Putative cold-tolerant genes identified in *S. cerevisiae* cause different phenotypes in the other *Saccharomyces* species

A previous study using a thermodynamic model combine with a large-scale competition experiment identified a list of candidate genes important for the growth at low temperature, among which *ADH3, GUT2, NMA1*, *YND1, FAA1*, and *ADH5*. Four out of six genes are involved in the cell redox balance: *i. ADH3, ADH5,* and *GUT2,* through their respective metabolic reactions, and *ii. NMA1* via NAD+ biosynthesis pathway (Fig. 2A) [35,36] *YND1* is involved in protein glycosylation, while *FAA1* is involved in long-chain fatty acid metabolism and import and hence may influence the transition of the cellular membrane to a more fluid state as temperatures decrease.

**Figure 2.**
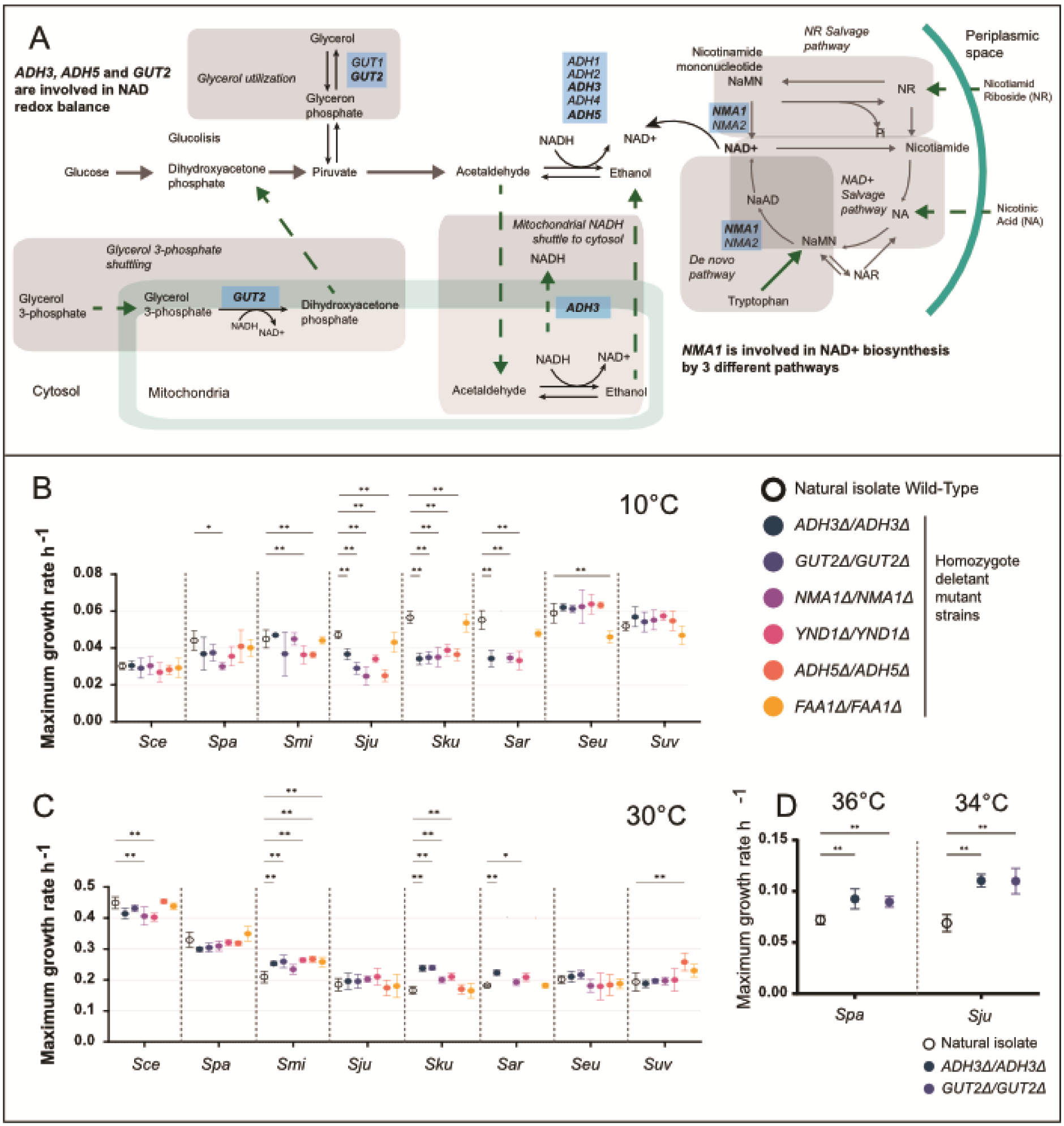
Maximum growth rate of the wild-type isolates and the homozygote deletant strains in eight species of the *Saccharomyces* genus. Metabolic pathways of *ADH3, ADH5* and *GUT2*. Pathways for NAD^+^ biosynthesis. Including three regulation pathways: de novo pathway (from tryptophan), NR salvage and NAD^+^ Salvage, by *NMA1* and its paralog gene *NMA2* (Panel A). The growth of all isolates was scored at 10°C (Panel B) and 30°C (Panel C). *S. paradoxus* and *S. jurei* natural isolates and the *ΔADH3* and *ΔGUT2* mutants was also scored at 36°C and 34°C, respectively (Panel D). Independents t-tests were performed pairing deletant mutants with natural isolates wild-types, p-values show significance at: *0.05, **0.01. *S. arboricola NMA1* and *ADH5* deletant mutants were excluded from the analysis.

Phenotypic effect of the deletions of *ADH3* and *GUT2* were previously validated in two natural species, namely S*. cerevisiae* and *S. kudriavzevii* [18].

Here, we systematically deleted the six candidate genes in all eight *Saccharomyces* species to understand their impact at genus level in conferring resistance to low temperatures. We observed that each deletion had a varying phenotypic effect according to the genetic background where it was introduced, and this was true both at low and high temperatures (Fig. 2). Strikingly, in *S. kudriavzevii*, *S. arboricola* and *S. jurei*, all candidate gene deletions had a large and significant impact on fitness at cold temperatures, with the exception of *FAA1*. Interestingly, the opposite behaviour was observed at warm temperature in *S. kudriavzevii* (but not in *S. jurei* or *S. arboricola*) where the deletion of four out of six genes led to improved fitness (Fig. 2C). In a similar fashion, for *S. mikatae*, deletion of *YND1* and *ADH5* cause a significant decrease in fitness at cold temperature (Fig. 2B), but, at 30°C, these same gene deletions, resulted in a significant fitness improvement (Fig. 2C).

While the deletion of *FAA1* at cold temperatures does not seem to affect *S. cerevisiae*, *S. paradoxus*, *S. mikatae*, *S. kudriavzevii*, *S. arboricola* and *S. jurei*; for *S. eubayanus*, this gene is the only one that affect significantly the fitness at cold. Although in *S. uvarum* no significant fitness changes were observed for any deletions tested, on average the deletion of *FAA1* gene also caused the bigger fitness impairment at cold (Fig. 2B).

At 30°C, the *ADH5Δ* mutant showed an improved fitness in *S. uvarum* and the *ADH3Δ* mutant displayed a growth advantage in *S. arboricola*. In *S. paradoxus*, only the *NMA1* deletion produced fitness defects at 10°C. Finally, in *S. cerevisiae* no phenotypic change was detected for any deletants tested at cold, however a small but significant fitness change was detected for *NMA1* and *YND1* at 30°C (Fig. 2C).

Taken altogether, these data suggest that the candidate genes tested play a crucial role in cold adaptation in a species-specific manner, and provide a further validation of the thermodynamic model prediction and genomic competition data our previous study [18].

In *S. paradoxus* and *S. jurei*, the largely unchanged fitness of the deletion mutants at 30°C may be due to the fact that this temperature is very close to their respective optimal growth conditions (Fig. 1). To investigate further this hypothesis, experiments were conducted at higher temperatures for these two species for *ADH3* and *GUT2*. Intriguingly, in both *S. paradoxus* and *S. jurei,* the deletion mutants exhibited a fitness improvement at 36°C and 34°C, respectively (Fig. 2D), suggesting that the effects of gene deletions on fitness become apparent at temperatures further from the species’ optimal range.

### Importance of transcriptional compensation in cold-tolerance response

*NMA1* is involved on the synthesis of NAD^+^, and its deletion partially reduces the capacity of the cell to synthetise NAD^+^ by the NAD^+^ salvage and nicotinamide riboside salvage pathways [36], affecting cellular redox reactions and the regulation of energy metabolism in the cell [37] (Fig. 2A).

The deletion of *NMA1* (nicotinamide mononucleotide adenylyl-transferase) revealed a fitness impairment in *S. paradoxus*, *S. jurei*, *S. kudriavzevii* and *S. arboricola* at cold temperature. No change in fitness was observed in species such as *S. cerevisiae, S. mikatae, S. eubayanus*, and *S. uvarum* upon *NMA1* deletion (Fig. 2B). Given that *NMA1* has a paralog, *NMA2*, we hypothesised that there may be an expressional compensation involving *NMA2* in the strain *NMA1Δ*. We assessed the mRNA levels of *NMA2*, at 10°C and al 30°C in all the species of the *Saccharomyces* genus (Fig. 3). The expression of *NMA2* at 10°C in *S. cerevisiae, S. eubayanus*, *S. uvarum*, was significantly higher in the *NMA1Δ* mutant strains compared to their respective wild-type strains, where *NMA2* expression was hardly detectable (Fig. 3A). In fact, it appears that the cells switch on the *NMA2* paralog upon deletion of *NMA1*. In *S. arboricola* we also detected an increase of *NMA2* expression in *NMA1Δ* strain, however here *NMA2* was also expressed in the WT (Fig. 3A). In the other species, the *NMA2* expression either did not change or decreased in the *NMA1* mutants (Fig. 3A). Besides being species-dependent, this transcriptional compensation also differs according to the temperature of growth. At 30°C, *NMA2* is expressed in all the WT species and corresponding *NMA1Δ* strains, but is increased only in *NMA1Δ* in *S. eubayanus* and *S. arboricola* background (Fig. 3B). Overall, this finding suggests that transcriptional activation of *NMA2* may compensate for the loss of *NMA1*, explaining the lack of fitness reduction at cold of *NMA1Δ* strains in *S. cerevisiae, S. eubayanus*, and *S. uvarum* (Fig. 2A).

**Figure 3.**
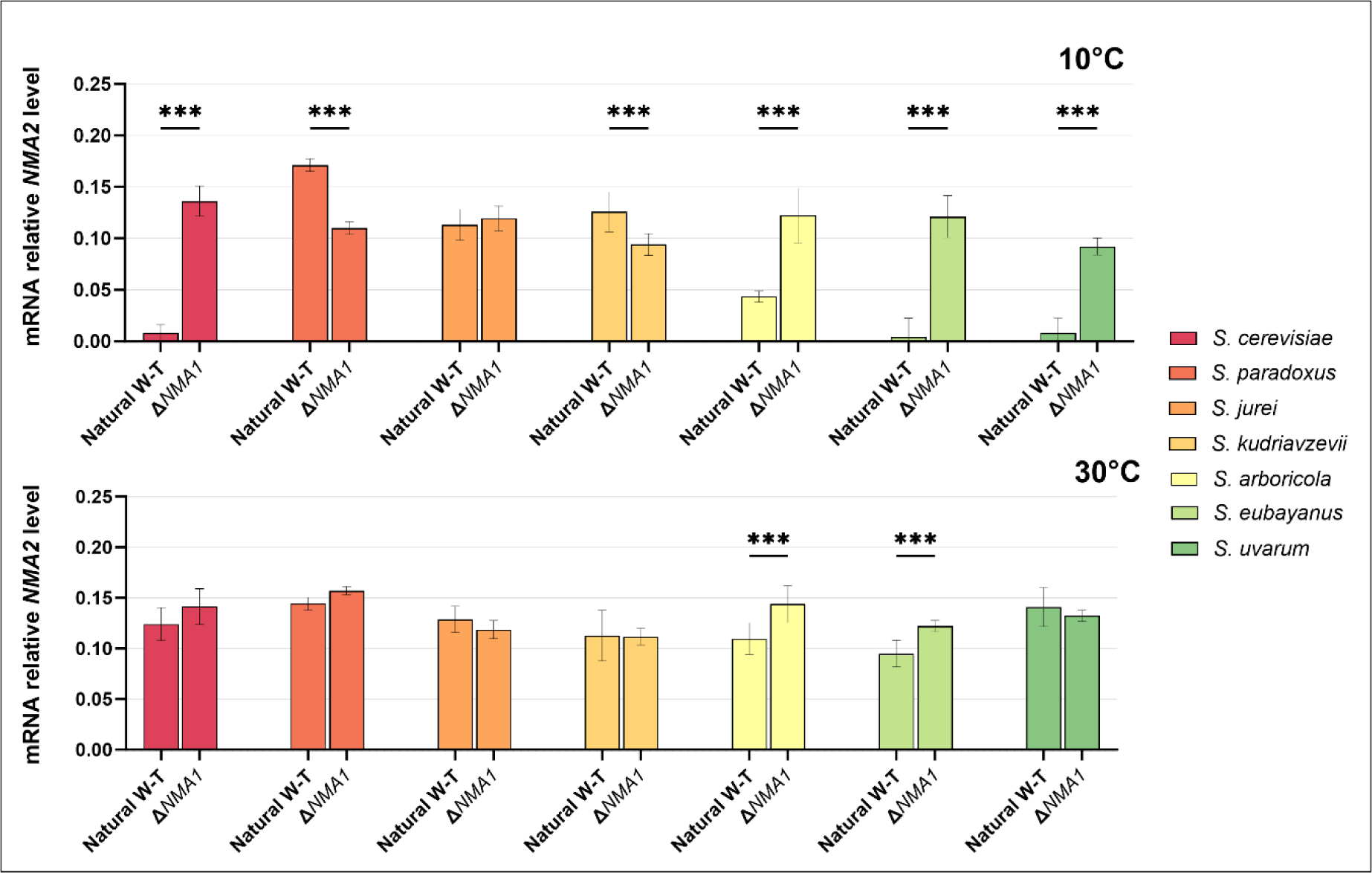
Relative mRNA levels of *NMA2* analysed by qRT-PCR in the natural yeast species and their respective *ΔNMA1* strains at 10°C and 30°C. p-values show significance at: *0.05, **0.01, and ***0.001.

### Identification of genetic interactions between candidate genes in the species of the *Saccharomyces* genus

We investigated the phenotypes of double mutants for all pair-wise combinations of five candidate genes (*ADH3*Δ*, GUT2*Δ*, NMA1*Δ*, YND1*Δ, and *FAA1*Δ) in five *Saccharomyces* species (*S. cerevisiae*, *S. paradoxus*, *S. jurei*, *S. kudriavzevii* and *S. uvarum*) to identify potential genetic interactions, either negative or positive, where the fitness of the double mutants is respectively lower or higher than the expected one based on the sum of the phenotypes of the single mutants.

We obtained all the double mutants with exception for Δ*GUT2/*Δ*YND1* in *S. jurei* for which transformation was not successful after several attempts. In total we created 49 double deletant mutants listed in Supplementary Table S1.

Genetic interactions for different candidate genes were scored at different temperatures, 10°C and 30°C. The presence/absence of interactions as well as the type of interactions, either negative or positive, was dependent on the growth temperature and the species background. The genetic interactions detected at warmer temperatures, were all positive in *S. jurei*, while in *S. kudriavzevii* and *S. uvarum* were all negative, involving different genes and only having one interaction in common (*YND1*/*ADH3*; Fig. 4). Interestingly in *S. uvarum*, *ADH3* showed interactions with all candidate genes except *FAA1*.

**Figure 4.**
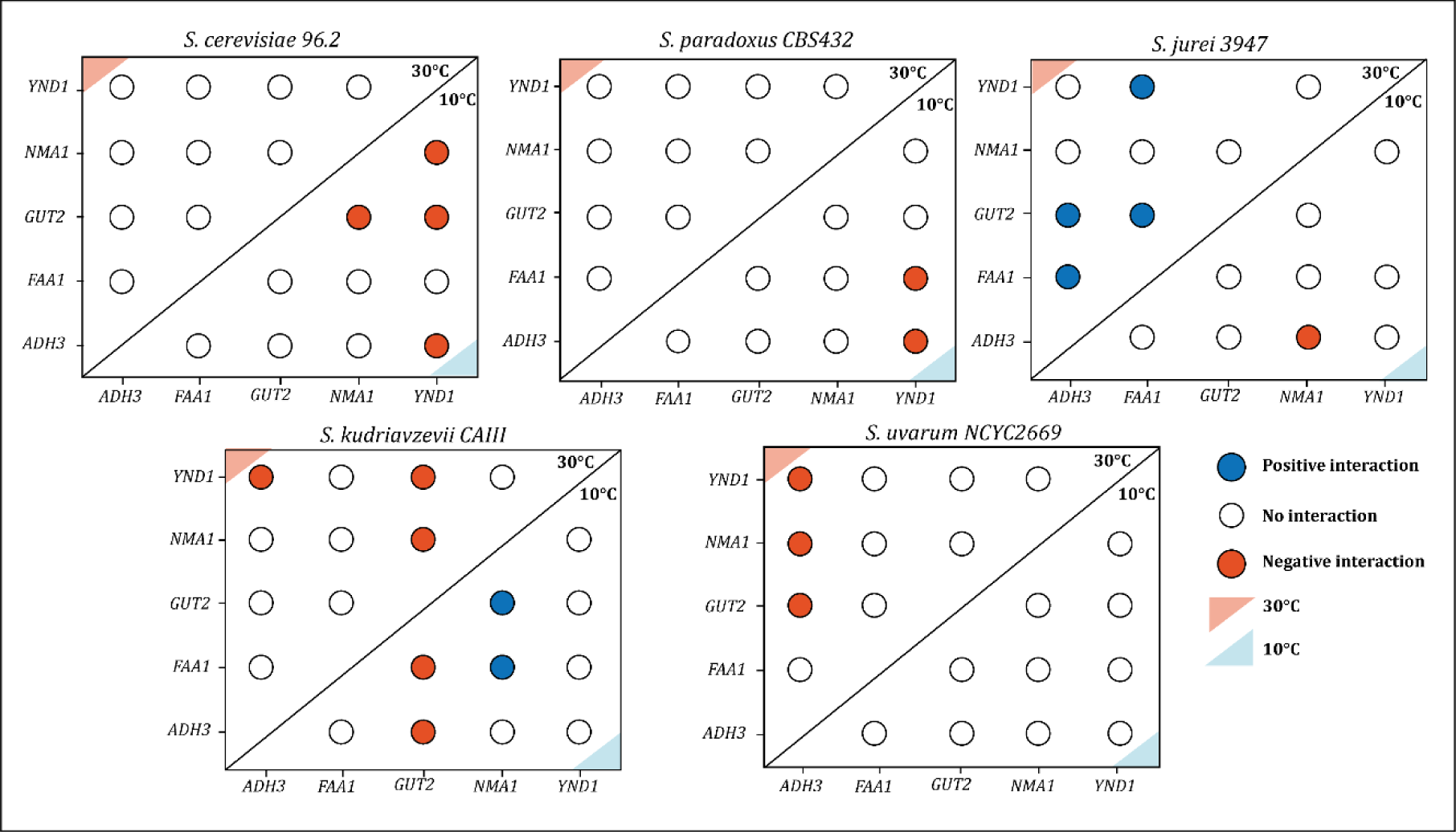
Absolute scores of genetic interactions in five candidate genes that provide resistance to cold in five species of the *Saccharomyces* genus. Blue and red coloured circles represent positive and negative interactions, respectively, calculated with an absolute genetic interaction score of |ε|>0.14 and a p value <0.05. Open white circles indicate no interactions. The upper left (light red triangle) and the lower right (light blue triangle) quadrants reports interactions at 30°C and at 10°C, respectively.

At cold temperature, all genetic interactions detected are negative for *S. cerevisiae*, *S. paradoxus* and *S. jurei*. No interactions were observed in *S. uvarum,* and *S. kudriavzevii* showed two interactions which were negative and two that were positive. The positive interactions in *S. kudriavzevii* involves the *NMA1* gene (Fig. 4).

In *S. cerevisiae* and *S. paradoxus*, four and two interactions, respectively, changed status from no interaction to negative when temperature dropped from at 30°C to 10°C (Fig. 4). Disruption of individual candidate genes did not affect fitness at cold temperatures in these species, but the combination of gene deletions hindered normal cellular development, resulting in lower fitness (negative interactions). In *S. jurei*, when shifting cells from 30°C to 10°C, we observed four interactions that change status (*i.e.* the double mutants *ΔADH3/ΔNMA1* that changes from no interaction at warm to negative at cold). Similarly, in *S. uvarum* and *S. kudriavzevii* the status of the interactions changes between warm and cold condition. Specifically, in *S. kudriavzevii* the double deletion *GUT2*/*NMA1* change directionality from positive at 30°C to negative at 10°C (Fig. 4). This double deletion has also a different impact on growth in *S. cerevisiae* and *S. kudriavzevii* background at cold.

This contrasting trend for the nature of interactions at different temperatures aligns well with the phenotypic differences scored between the thermo-tolerant *S. cerevisiae* and the cold-tolerant *S. kudriavzevii* (Fig. 2B and 2C).

Additionally, we investigated the interactions of between *ADH3*, *GUT2* and *NMA1* and two intergenic non-coding RNA (ncRNA), SUT125 and SUT035, which were identified as important for growth at low temperature and for transcription of genes involved in mitochondrial functions [26,27]. We created the double deletions (six in total) in *S. cerevisiae* background. We were able to observe several genetic interactions which again appeared to be temperature-dependent (Fig. S1). In particular, significant negative interactions were scored between SUT035 and all the gene tested at 30°C. This exploratory data adds evidence that the mechanism behind temperature adaptation is not solely reliant on protein-coding genes. The strains used are listed in Supplementary Table S1.

### Native transcription of candidate genes reveals different strengths of gene expression at cold in the species of *Saccharomyces* genus

We next sought to determine the strength of the expression of the candidate genes at cold in the different species to determine whether it may correlate with the phenotype observed upon their deletion. In fact, expressional fluctuations could play a role in cold-tolerance alongside the type of alleles [8,38].

Firstly, we compared the mRNA levels of *ADH3*, *GUT2*, *NMA1*, *YND1*, *ADH5* and *FAA1* at 10°C and 30°C in all the species belonging to the *Saccharomyces* genus. Our results revealed distinct species-specific expression patterns for the candidate genes, highlighting the complexity of transcriptional regulation within the *Saccharomyces* genus. *FAA1* displayed a relatively consistent expression pattern across species and temperatures. Despite potential disruptions in the synthesis and activation of fatty acids caused by temperature changes, the activation of exogenous fatty acids mediated by *FAA1* appears to remain constant at both warm and cold temperatures, with consistent expression observed across all *Saccharomyces* species [39,40,41].

Upon analysing the expression of the candidate genes at 10°C, we observed that *ADH3*, *NMA1*, *YND1, GUT2* and *ADH5* showed higher expression in *S. kudriavzevii, S. jurei*, *S. mikatae* and *S. paradoxus*, compared to the other species. In contrast*, S. cerevisiae*, *S. arboricola*, *S. eubayanus* and *S. uvarum* showed the low mRNA levels for all the above-mentioned genes (Fig. 5). These results explain also the phenotypic data for *S. eubayanus* and *S. uvarum* (Fig. 2B) where the deletion of all the redox genes do not impair phenotype at cold, likely because from the start they are poorly expressed in the wild-type. Additionally, we measured to expression of the candidate genes at 30°C (Fig. S2).

**Figure 5.**
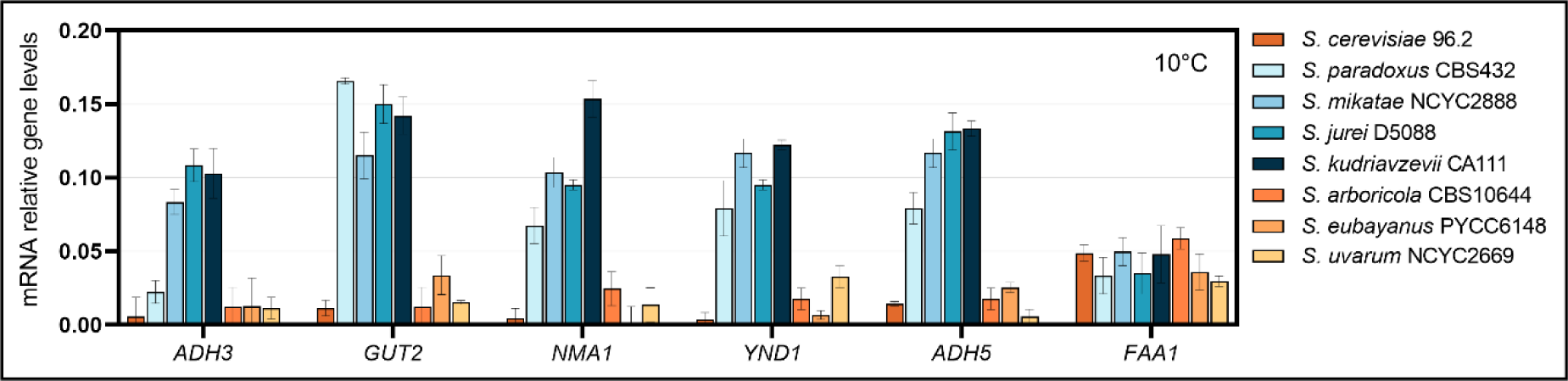
Relative mRNA levels of *ADH3*, *GUT2*, *NMA1*, *YND1*, *ADH5*, and *FAA1* analysed by qPCR in natural isolated strains of eight species of the *Saccharomyces* genus.

### *S. kudriavzevii* promoter and allele swap experiments support the role of the expression over the allele type in cryotolerance

Since we observed a strong expression of the candidate genes in the natural cold-tolerant *S. kudriavzevii* CAIII strain, we sought to investigate whether is the promoter (transcription) or the allele (genetic) the main player (factor) in the cold tolerance trait. First, we ran a SIFT analysis on all the alleles of the six candidate genes to see how many amino acids changes were present between *S. cerevisiae* and *S. kudriavzevii* that can potentially affect the protein structure (Supplementary Table S2). Changes that affected conserved regions were detected, with the *YND1* allele harbouring the highest number of them (7 amino acid changes; Table S2) and no changes in *ADH3, NMA1* and *FAA1*. Based on the SIFT analysis, we selected *YND1* and *ADH3* (predicted to have both conserved region and protein stability affected) as candidates to evaluate the relevance of the allele sequence vs the promoter in cold adaptation.

The effect of the promoter and allele on the growth phenotype of the different *Saccharomyces* species was investigated by promoter and allele replacement experiments.

For the promoter swap, plasmids were constructed (Table S4) with the *S. kudriavzevii* promoter (*Pk*) placed in front of *S. cerevisiae, S. paradoxus, S. jurei and S. eubayanus ADH3* and *YND1* alleles. For the allele swap, plasmids were constructed each containing the native species promoter in front of the *S. kudriavzevii* allele (*Ak*) of *ADH3* and *YND1* genes. As control, plasmids with the different native promoters and native alleles were generated (Supplementary Fig. S3). All the plasmids were then introduced in the relevant homozygote deletant *ADH3D* or *YND1D* strains to score the phenotype.

The fitness of *S. cerevisiae, S. paradoxus, S. jurei and S. eubayanus* carrying *Pk-ADH3, Pk-YND1, Ak-ADH3* and *Ak-YND1* were scored in liquid media at 10°C and 30°C. The fitness score of the strains carrying the constructed plasmid was inferred from the integral area under the curve and compared to the fitness of the natural W-T to obtain the growth ratio.

The promoter replacement experiment revealed an increment of fitness at 10°C in all strains carrying the promoter of *S. kudriavzevii ADH3* (Table 1A) and *YND1* (Table 2A), while the controls with their natural promoter showed non-significant differences in fitness compared to the wild-type strain. At 30°C, no significant changes were observed on the strains carrying the promoter, except for *S. jurei* where a fitness improvement for *ADH3* for the strain carrying the *S. kudriavzevii* promoter was observed (Table 1A; Table 2A).

**Table 1.** Growth ratio of yeast cells containing prs418 plasmid with either A) Pk-*ADH3* + natural *ADH3* allele, or B) natural *ADH3* promoter + Ak-*ADH3*. p-values show significance at: *0.05, **0.01, ns= no significant.

|  |
| --- |
| **A** |
| *Integration of prs418 plasmid with S. kudriavzevii ADH3 promoter* |

| strain + *S. kudriavzevii*<br>*ADH3* promoter | Ratio to wild<br>type at 10°C | p value | Ratio to wild<br>type at 30°C | p value |
| --- | --- | --- | --- | --- |
| *S. cerevisiae* | **1.34** | ** | 0.99 | ns |
| *S. paradoxus* | **1.39** | ** | 0.81 | ns |
| *S. jurei* | **1.38** | ** | **1.66** | ** |
| *S. eubayanus* | **1.38** | ** | 0.96 | ns |
| Control *S. cerevisiae* | 1.04 | ns | 0.98 | ns |
| Control *S. paradoxus* | 0.98 | ns | 0.96 | ns |
| Control *S. jurei* | 1.00 | ns | 1.12 | ns |
| Control *S. eubayanus* | 0.92 | ns | 0.94 | ns |
| **B** |  |  |  |  |
| *Integration of prs418 plasmid with S. kudriavzevii ADH3 allele* |  |  |  |  |
| strain + *S. kudriavzevii*<br>*ADH3* allele | Ratio to wild<br>type at 10°C | p value | Ratio to wild<br>type at 30°C | p value |
| *S. cerevisiae* | **0.79** | ** | 0.97 | ns |
| *S. paradoxus* | **0.82** | ** | 1.01 | ns |
| *S. jurei* | **0.81** | ** | 0.93 | ns |
| *S. eubayanus* | **0.89** | ** | 0.99 | ns |
| Control *S. cerevisiae* | 0.97 | ns | 0.96 | ns |
| Control *S. eubayanus* | 0.99 | ns | 0.99 | ns |
| Control *S. jurei* | 0.92 | ns | **1.18** | * |
| Control *S. eubayanus* | 0.93 | ns | 1.01 | ns |

**Table 2.** Growth ratio of yeast cells containing prs418 plasmid with either A) Pk-*YND1* + natural *YND1* allele, or B) natural *YND1* promoter + Ak-*YND1*. p-values show significance at: *0.05, **0.01, ns= no significant.

| **A** |  |  |  |  |
| --- | --- | --- | --- | --- |
| *Integration of prs418 plasmid with S. kudriavzevii YND1 promoter* |  |  |  |  |
| strain + *S. kudriavzevii*<br>*YND1* promoter | Ratio to wild<br>type at 10°C | p value | Ratio to wild<br>type at 30°C | p value |
| *S. cerevisiae* | **1.29** | ** | 1.02 | ns |
| *S. paradoxus* | **1.38** | ** | 0.91 | ns |
| *S. jurei* | **1.38** | ** | 0.93 | ns |
| *S. eubayanus* | **1.22** | ** | 0.99 | ns |
| Control *S. cerevisiae* | 0.99 | ns | 0.93 | ns |
| Control *S. paradoxus* | 1.01 | ns | 0.99 | ns |
| Control *S. jurei* | 1.09 | ns | 0.98 | ns |
| Control *S. eubayanus* | 0.97 | ns | 1.00 | ns |
| **B** |  |  |  |  |
| *Integration of prs418 plasmid with S. kudriavzevii YND1 allele* |  |  |  |  |
| strain + *S. kudriavzevii*<br>*YND1* allele | Ratio to wild<br>type at 10°C | p value | Ratio to wild<br>type at 30°C | p value |
| *S. cerevisiae* | 0.98 | ns | 1.18 | ns |
| *S. paradoxus* | **0.82** | ** | 0.95 | ns |
| *S. jurei* | 0.98 | ns | 0.95 | ns |
| *S. eubayanus* | 0.93 | ns | 1.09 | ns |
| Control *S. cerevisiae* | 0.92 | ns | 1.01 | ns |
| Control *S. eubayanus* | 0.99 | ns | 0.96 | ns |
| Control *S. jurei* | 1.08 | ns | 0.99 | ns |
| Control *S. eubayanus* | 1.11 | ns | 1.00 | ns |

Replacing the *ADH3* natural allele with the *S. kudriavzevii* allele (*Ak-ADH3*) in *S. cerevisiae*, *S. paradoxus*, *S. jurei* and *S. eubayanus* produced a significant reduction in fitness at 10°C compared with the natural isolate, while no significant changes were seen at 30°C for any strains tested (Table 1B). Thus, even if the similarity of the *S. kudriavzevii ADH3* sequence is ≥97% compared to the native alleles we can conclude that the insertion of a heterologous allele is not optimal and may interfere on protein-protein interactions causing sub-optimal growth (Cherry et al., 1998; Scannell et al., 2011).

The replacement of the *S. kudriavzevii YND1* allele (*Ak-YND1*) resulted in non-significant changes in fitness at 10°C or 30°C, except for *S. paradoxus* at 10°C (Table 2B).

The results of the promoter and allele replacement suggest that the role of the promoter is more important than the allele to provide resistance to cold in *Saccharomyces* species. One explanation is the up-regulation of the alcohol dehydrogenase-3 produced in the mitochondria due to the Pk*-ADH3* increase the fitness of the yeast species at cold but not at warm temperatures. The ethanol-acetaldehyde shuttle between the mitochondrial matrix and the cytosol may increase due *ADH3* up-regulation, maintaining the mitochondrial redox balance at low temperatures [18,28].

We next checked if the Pk does indeed increase the expression of the downstream genes. Changes of *ADH3* gene expression in *S. cerevisiae, S. paradoxus, S. jurei, S. eubayanus* and *S. kudriavzevii* carrying the cold-tolerant *S. kudriavzevii* promoter were scored to identify the effect of the *S. kudriavzevii* promoter on *ADH3* expression in different *Saccharomyces* species at 10°C and 30°C. We confirmed that *pK* enhanced the expression of the *ADH3* gene in *S. cerevisiae S. paradoxus, S. jurei and S. eubayanus* at both 10°C and 30°C temperatures, confirming that the expression of alcohol dehydrogenase-3 enzyme is higher under Pk than under the natural promoters in *S. cerevisiae*, *S. paradoxus*, *S. jurei* and *S. eubayanus* (Supplementary Fig. S4). Additionally, in *S. cerevisiae* we also checked the expression of *YND1* under the *Pk*, and again we observed an increase of expression (Supplementary Fig. S4).

In conclusion, this experimental approach was sufficient to identify phenotypic differences due to regulation *in cis* [42]. We confirmed that *S. kudriavzevii* promoter has a stronger effect upon gene expression and phenotypes. Therefore, transcriptional adaptation may have occurred within the *Saccharomyces* genus to allow yeast species to adapt to new environmental conditions by constantly modifying the expression of their genes [8], allowing them to co-exist by occupying the same habitat but using different thermic niches. These results open the door for further investigation on transcriptional adaptation of species under fluctuating temperatures.

## Discussion

In this study, different approaches were used to uncover the effect of cold temperature on yeast species belonging to *Saccharomyces* genus and to understand the molecular mechanisms that affect growth at cold temperatures, enabling a better understanding and how biodiversity originates and is maintained in constantly changing environments. The data obtained in this study suggests that cold temperature is a key factor that allows the species of the *Saccharomyces* genus to diversify and potentially occupy the same niche [13,34,43].

The disruption of candidate genes responsible for adaptation to cold allowed us to evaluate its importance in a multi-species background. The results suggested that putative cold-tolerant genes identified in *S. cerevisiae* cause diverse phenotypes in the other *Saccharomyces* species. The deletion of *ADH3* and *GUT2,* both in homozygosis and heterozygosis, has already been showed to impair fitness in the cold-tolerant *S. kudriavzevii* at 12°C [18] in chemically defined media limited either for carbon or nitrogen.

The genes *ADH3, ADH5, NMA1 YND1 and GUT2* help to maintain redox balance between NADH and NAD^+^, causing a conservation of NADH or favouring either ethanol or glycerol production in the case of *ADH3* and *GUT2*, respectively (44, 45). High expression of alcohol dehydrogenase genes (*ADH*) has been observed as a response to cold adaptation in various organism. For instance, the extremophilic yeast *Rhodotorula frigidialcoholis* overexpressed *ADH* at 0°C [46]. A similar upregulation of *ADH* was also observed in the Arctic permafrost bacterium, *Planococcus halocryophilus*, when grown at −15 °C [47]. In the context of yeast ethanol production, *S. cerevisiae* has been reported to produce ethanol at low temperatures during wine fermentation (0 and 2 °C), although these studies involved the addition of biocatalysts to facilitate fermentation [48].

*YND1* is involved in protein glycosylation, while *FAA1* is involved in long-chain fatty acid metabolism and import and hence may influence the transition of the cellular membrane to a more fluid state as temperatures decrease.

Glycerol is known as an effective cryoprotectant for yeast, hindering the hydrogen bonding in water molecules [44]. The significance of intracellular glycerol as a cryoprotectant is evident in the cold-tolerant *S. kudriavzevii*, which produces higher glycerol concentrations compared to the thermotolerant *S. cerevisiae* [49]. Therefore, the reduction in fitness in *S. kudriavzevii* observed at cold temperatures upon *GUT2* deletion could be attributed to the importance of intracellular glycerol production for cryoprotection. In contrast, *FAA1*, a gene associated with membrane fluidity, appeared to have a predominant role in cold adaptation for *S. eubayanus* and *S. uvarum*. Cold survival of cells relies on their ability to adjust membrane composition to maintain fluidity and avoid a gel-like state, which could be detrimental [41,50]. *Faa1p* catalyses the activation of long fatty acids ranging from 12 to 16 carbons and has the main acyl-CoA synthetase activity within the cell [30,51]. Thus, at cold temperatures, *FAA1* is upregulated to speed up the membrane biogenesis towards a more fluid state.

The role of *ADH3* and *GUT2* in maintaining mitochondrial redox balance and glycerol metabolism, respectively, sheds light on the observed fitness effects upon their deletion. *ADH3* is involved in the ethanol-acetaldehyde shuttle, which helps maintain mitochondrial redox balance by facilitating the oxidation of NADH in the cytosol [28]. On the other hand, *GUT2* plays a role in the glycerol-3-phosphate shuttle within the mitochondria [31]. Conversely, the fitness improvement upon deletion of *ADH3* and *GUT2* at warm temperatures in *S. mikatae, S. kudriavzevii and S. arboricola* could be attributed to several factors. The deletion of *ADH3* may result in an increment of acetaldehyde concentration and activity in the cytosol leading to a redox imbalance that can be reversed by increasing glycerol production [13,18,28] (Fig. 2A). Additionally, the fitness advantage upon *GUT2* deletion at 30°C could be linked to the upregulation of glycerol active transporters triggered by the temperature increase [52]. These mechanisms contribute to the observed fitness improvements in the mentioned species at higher temperatures.

The deletion of *YND1*, a gene that encodes an apyrase involved in protein traffic instead of redox balance, produces a reduction of the fitness at cold in *S. mikatae*. *S. jurei*, *S. kudriavzevii* and *S. arboricola*, probably due to decreased glycosylation in the cell [53]. *YND1* is also involved in the membrane compartmentalization by sphingolipids, to shape microdomains within the membrane, and their synthesis has been reported to be related to high temperature related stress in *S. cerevisiae* [54, 55, 56]. Due to an increased rigidity of the plasma membrane caused by low temperature, an impairment of tryptophan transport may occur. The NAD^+^ biosynthesis in the *de novo* pathway may be affected (Fig. 2A), increasing the activity in the Salvage pathways. The deletion of *NMA1* (Nicotinamide Mononucleotide Adenylyl-transferase) partially reduces the capacity of the cell to synthetise NAD by the NAD^+^ salvage and nicotinamide riboside salvage pathways [36], hindering cellular redox reactions and energy metabolism in cold-tolerant species. Given that NAD^+^ is involved in the regulation of energy metabolism, the disruption of genes involved in NAD^+^ redox balance may be affected, beside metabolic pathways, several other biological processes, including DNA repair and transcription may also be impaired [36,37].

It is known that the two cryo-tolerant species *S. kudriavzevii* and *S. uvarum* have developed different strategies for cold resistance. Pathways in production of NAD^+^ play a major role in cold adaptation in *S. kudriavzevii* while changes in the biosynthesis of folates and aromatic amino acids pathway (Shikimate) plays a significant role in *S. uvarum* [57]. Our data shows that *S. uvarum* with *S. eubayanus* have a similar phenotypic profile upon *FAA1* deletion therefore is possible that both species use the same strategy for cold tolerance. This would also resonate with the fact that *S. uvarum* and *S. eubayanus* are phylogenetically closely related.

Previous literature has described examples of negative epistasis between unlinked adaptive genes during evolution experiments [58,59]. Positive epistasis interactions may lead to mutations that cause big fitness changes, speeding up adaptation [60]. The positive epistasis between genes or between genes and ncRNAs support this type of evolutionary dynamics. Epistatic interactions between candidate genes are revealed in all species but appear to be temperature dependant. As mentioned previously, it is known that cold adaptation strategies vary between species, therefore, the types of genetic interactions between candidate genes can vary depending on the species cold adaptation strategy.

After the replacement of *S. cerevisiae, S. paradoxus, S. jurei and S. eubayanus ADH3* and *YND1* promoters with *S. kudriavzevii* promoters, we identified that the role of the promoter is more important than the allele type to provide optimal growth at colder temperatures. One explanation for the higher fitness of the strains containing the *S. kudriavzevii ADH3* promoter swap in *S. cerevisiae*, *S. paradoxus*, *S. jurei* and *S. eubayanus*, is that the alcohol dehydrogenase-3 mobilised in the mitochondria is upregulated by recruiting more effectively transcription factors in the *S. kudriavzevii ADH3* promoter, increasing the fitness at cold. In the case of *S. kudriavzevii YND1* promoter integration in the same set of species, sphingolipid synthesis may be upregulated by the in-cis transcription factors, also triggered by cold temperature, changing the membrane compartmentalization which is mediated by sphingolipids. These lipids shape microdomains within the membrane, and their synthesis has been reported to be related to high temperature related stress in *S. cerevisiae* [54,55].

## Conclusions

This study provides valuable insights into the molecular factors influencing temperature-dependent growth profiles *Saccharomyces species*, shedding light on the importance of cold-tolerance in the diversification and adaptation of these yeasts. A number of genes, namely *ADH3, GUT2, NMA1, YND1, FAA1*, and *ADH5*, were identified as crucial players in cold adaptation, involving either in the conservation of redox-balance or in the long-chain fatty acid metabolism. We investigated genetic interactions in multispecies background, identified cases of environmental plasticity of epistasis, and highlighted the diverse strategies employed by different species to adapt to varying temperatures. Our findings also reveal the complexity of transcriptional regulation within the *Saccharomyces* genus. Promoter swap experiments demonstrated that the *S. kudriavzevii* promoter enhanced expression of *ADH3* and *YND1* in other *Saccharomyces species* and drove fitness improvement at low temperatures. These data underscore the phenotypic impact of transcription over allele type, supporting the notion that stochastic tuning of the expression network may have driven temperature adaptation.

In conclusion, by unravelling the interplay between gene expression, allele variation, genetic background and environment, this study emphasizes the intricate nature of transcriptional regulation and its pivotal role in facilitating cold adaptation in *Saccharomyces* species.

## Methods

### Strains and plasmids

The strains used un this study are natural isolates wild-types: *S. cerevisiae S.C96.2*, *S. paradoxus CBS432, S. mikatae NCYC2888, S. jurei D5088, S. kudriavzevii CA111, S. arboricola CBS10644, S. eubayanus PYCC6148, S. uvarum NCYC2669.* All the *Saccharomyces* strains provided and constructed in this work are listed in Supplementary Table S1. All the plasmid used and constructed in this study are reported in Supplementary Table S4. Briefly, the pUG-6 plasmid was used to amplify the *loxP-kanMX-loxP* cassette, while pRS418 (Addgene plasmid #11256) was used to amplify the *clonNAT* cassette and also was the vector of choice for Gibson assembly cloning methodology for the promoter/allele swap experiments.

### Fitness assays

Fitness assays were performed using a plate reader from FLUOstar OPTIMA plate reader (BMG Labtech, UK). To obtain the growth curves we inoculate cells to an OD_600nm_ = 0.1 which equals ∼10^6^ cells in 200 μL of YPD media (20g/L peptone, 10g/L yeast extract, 2% glucose). We measured the optical density of the samples every 5 minutes for 24 hours for the cells growing at 30°C and for 72 hours for the ones growing at 10°C. Blank-corrected data was used to obtain fitness scores in terms of maximum growth rate, maximum biomass and are area under growing curve, using *gcFit* and *gcPlot* function included in the groFit R package [61]. To obtain optimal growth temperature range, maximum growth rates scores were obtained for temperatures ranging from cold temperature (10°C) with steps of 5°C until warm temperature, 40°C. The optimal growth temperature for each species was estimated using a non-linear model [13,62] in this case we used a third order polynomial curve fitting using GraphPad Prism version 9 software.

### Creation of gene deletion mutants

The genes: *ADH3, GUT2, NMA1, YND1, ADH5* and *FAA1* were deleted by the insertion of *loxP-kanMX-loxP* [63] cassette into the cell via homologous recombination using 45 bp 3’ and 5’ overhang homology sequences designed for each *Saccharomyces* species, which enabled species-specific targeting of the gene (Supplementary Table S3). In total 50 deletion cassettes were amplified via PCR and the cassettes were inserted into the genome using Li/Ac transformation protocol [64].

#### Homozygous mutant line generation

The creation of homozygote deletion mutants was achieved by sporulation of the diploid heterozygote transformants using potassium acetate media plates, which triggered meiosis due to nitrogen and carbon starvation. Spores were observed after 4-7 days and were dissected through digestion of the ascospore wall within a digestion solution (5ng/μL lyticase in 1.5M Sorbitol) and incubating the solution at 37°C for 10 minutes. The tetrads were separated/dissected using a Singer micromanipulator (Singer instruments, UK) on YPAD-G418 (300 mg/L) plates. The homothallic strains result on a diploid colony by self-fertilization. A set of homozygotes deletant strains for *ADH3*, *NMA1*, *YND1, GUT2, ADH5* and *FAA1* genes were created in *S. cerevisiae*, *S. paradoxus*, *S. mikatae*, *S. jurei*, *S. kudriavzevii*, *S. eubayanus* and *S. uvarum* (Supplementary Table S1). In *S. arboricola,* mutants were created for *ADH3*, *NMA1*, *YND1* and *FAA1,* as gene deletion was unsuccessful for *GUT2* and *ADH5*.

### Creation of double deletant mutants

We created a combination of double gene deletant mutants on *S. cerevisiae, S. paradoxus, S. jurei, S. kudriavzevii* and *S. uvarum*. We used homozygote deletion mutants whose knock-out was achieved by the insertion *loxP-kanMX-loxP* cassette. An additional *loxP-cloNAT-loxP* [63] cassette was into the cell via homologous recombination to replace the ORF of the second gene of interest. Via sporulation and dissection of tetrads in plates containing both nourseothricin 100 (µg/ml) and neomycin (200 µg/ml) drugs, we obtained homozygote double deletant strains.

### Analysis of the genetic interactions

We obtained all the colony size of the mutants using the Phenobooth (Singer Instruments Ltd, Somerset, UK). The colony sizes of both double and single mutants were normalized per plate. The interaction score ε was calculated according to [65,66]. Briefly, ε was obtained by comparing the single mutant fitness (*i.e.,* WA, WB) to the double mutant fitness (WAB) as following

Score ε = WAB – WA x WB

The absolute genetic interaction score of |ε| > 0.14, and a p-value < 0.05 were used as a defined confidence threshold for significant interactions.

### RNA extraction and quantitative Real Time-PCR (qRT-PCR)

RNA was extracted from three biological replicas of cells in the mid-log phase of growth (OD_600nm_ = 0.4 – 0.6), growing at 10°C and 30°C in YPD media, using a RNeasy kit from QIAGEN. Quality and concentration or the RNA samples was assessed through spectrophotometry using a NanoDrop (Thermo Scientific) Furthermore, the integrity of the RNA samples was assessed by running the denatured samples in a 1% agarose gel in 1X TAE buffer (40mM Tris, 20mM acetic acid, 1mM EDTA) at 70V for 1 hour. cDNA was synthetized using a Tetro cDNA synthesis kit (Meridian). Quantitative Real-Time PCR was used to measure the relative gene expression of the candidate genes using three biological replicates and three technical replicates per species and treatment. Real-time PCR was run in a Lightcycler 480 (Roche), using SYBR Green Master Mix–BioRad as a fluorescent dye. The oligos were designed to amplify 200-350 bp with the gene of interest (Supplementary Table S3).

The data obtained in the qRT-PCR experiments was analysed manually as relative quantification by measuring *ΔCt* which is equal to the difference in the fluorescence detection above a certain threshold of the genes being compared [67]. In this case the reference gene used was *ACT1*. *ΔCt* of the candidate genes and the *ACT1* gene was calculated by subtracting the *Ct* of 3 technical replicates of the candidate gene to the average *Ct* of the *ACT1* gene. For each species, the resulting *ΔCts* values were averaged among the total nine replicates (*i.e.,* each of the 3 biological replicas had 3 technical replicas) and were normalized by calculating the logarithmic value for visualization purposes.

### Sorting intolerant from tolerant (SIFT) analysis

The SIFT methods using Mutfunc was used to predicts whether an *amino acid* substitution affects protein function based on sequence homology and the physical properties of *amino acids* [68]. Mutfunc is a resource used to annotate variants, displaying the ones that are likely to be deleterious to function and predicted consequences on protein stability, interaction interfaces, regulatory regions (TF binding sites), linear motifs and conservation. The annotations/predictions are based on the computation on the impact of all possible variants using existing algorithms that cover different mechanisms [69]. The number of amino acids changes and their potential effect on the proteins in *S. kudriavzevii*, was obtained using *S. cerevisiae* candidates genes sequences as reference.

### Promoter and allele swap

*S. cerevisiae, S. paradoxus, S. jurei and S. eubayanus ADH3* and *YND1* native promoters were replaced with “*S. kudriavzevii* promoter” (Pk). Additionally, in the same species the *ADH3* and *YND1* alleles were replaced with the “*S. kudriavzevii* allele” (Ak), placing them under the regulation of *S. cerevisiae, S. paradoxus, S. jurei and S. eubayanus* native promoters. Gibson assembly cloning protocol was employed to integrate DNA fragments in the linearized vector by mainly three reactions: 5′ exonucleases, the 3′-extension activity of a DNA polymerase and DNA ligase activity (Supplementary Fig. S5). A region of about 800 bp 5’ upstream of the ORF of the candidate genes was swapped to evaluate the importance of *in cis* transcription of genes at different temperatures. Two sets of five plasmids for each gene were constructed including *i.* the 4 host alleles under the Pk (promoter swap), and *ii*. the *Ak* under the four host promoters (allele swap). Given that two genes were investigated, a total of 20 plasmids were constructed, including control plasmids carrying concomitantly both native promoters and native alleles for each species background (Supplementary Table S4). The plasmids constructed were introduced into single homozygote deletant strains for *ADH3* and *YND1*, independently. Fitness assays were carried out in natural wild-types, allele and promoter swapped strains. Specific primers were used for subsequent Gibson assembly cloning methodology (Supplementary Table S3).

## Acknowledgements

We thank Gonzalo Gómez Cepa for his help with some deletion experiments.

## Authors contribution

JP: conceptualization, investigation, methodology, formal analysis, validation, visualization, writing-original draft, writing-review and editing. DD: project administration, supervision, conceptualization, formal analysis, writing-original draft, writing-review and editing. LNBC: methodology, formal analysis, writing-review and editing.

## Supporting information captions

*Supplementary figure S1*. Absolute scores of genetic interactions of *ADH3, GUT2* and *NMA1* with the ncRNA transcript SUT125 (A) and SUT035 (B) at 30°C and 10°C. Blues dots represents positive interactions; red dots, negative interactions; and greys dots, no interaction.

*Supplementary figure S2*. Relative mRNA levels of *ADH3*, *GUT2*, *NMA1*, *YND1*, *ADH5*, and *FAA1* analysed by qPCR in natural isolated strains of eight species of the *Saccharomyces* genus at 30°C.

*Supplementary Figure S3.* Strategy for the plasmid assembly carrying *ADH3* and *YND1 S. kudriavzevii* promoter with *ADH3* (A) and *YND1* (C) *Saccharomyces* species native allele, respectively; and the assembly of plasmids carrying *ADH3* and *YND1 Saccharomyces* species native promoters upstream *ADH3* (B) and *YND1* (D) *S. kudriavzevii* alleles.

*Supplementary figure S4.* Panel A: relative mRNA levels of *ADH3* in *S. cerevisiae, S. paradoxus, S. jurei* and *S. eubayanus* natural W-T and their respective mutants carrying *Pk-ADH3* (*S. kudriavzevii ADH3* promoter). Panel B: relative mRNA levels of *YND1* in *S. cerevisiae* natural W-T and their respective mutants carrying *Pk-YND1* (*S. kudriavzevii YND1* promoter). *p*-values are indicated as: \*\*\*\**P* < 0.0001.

*Supplementary figure S5*. Gibson Assembly technique to construct plasmids for *S. kudriavzevii* promoter and allele swap. (A) Diagram of the assembly steps: 5′ exonucleases, the 3′-extension activity of a DNA polymerase and DNA ligase activity, the diagram includes sizes of overlapping section between fragments and PCR melting temperature. (B) Agarose gel showing on the left the amplification band of prS418 plasmid (empty vector) and on the right the amplification bands of YND1 alleles and promoters of *S. cerevisiae* (Sc), *S. paradoxus* (Sp), *S. jurei* (Sj), *S. eubayanus* (Se) and *S. kudriavzevii* (Sk). Specific primers were used for subsequent Gibson Assembly cloning.

*Supplementary Table S1*. List of yeast strains used in this study

*Supplementary Table S2*. Number of amino acids changes and their potential effect on the proteins, obtained by SIFT analysis using S. cerevisiae sequences as reference. The total number (#) of amino acids changes refers to changes in S. kudriavzevii. *Site of amino acid change that affect protein function.

*Supplementary Table S3*. Oligos used to incorporate native S. kudriavzevii promoter + Saccharomyces allele into plasmid prs418 via Gibson Assembly

*Supplementary Table S4*. List of plasmids used in this study

